# Integrating longitudinal hyperspectral phenotyping with AI and GWAS to dissect barley waterlogging responses

**DOI:** 10.64898/2026.06.02.729597

**Authors:** Villő Bernád, Jason J. Walsh, Emilie Jacob, Mortaza Khodaeiaminjan, Nick Zhang, Keru Wang, Faezeh Fadaei, Laura Craig, Wagner Barreto-Souza, Eleni Mangina, Laurent Gutierrez, Sónia Negrão

**Affiliations:** School of Biology and Environmental Science, University College Dublin, Dublin, Ireland; Centre de Ressources Régionales en Biologie Moléculaire, Université de Picardie Jules Verne, Amiens, France; School of Mathematics and Statistics, University College Dublin, Dublin, Ireland; School of Computer Science, University College Dublin, Dublin, Ireland

**Keywords:** Artificial intelligence, Barley, GWAS, High-throughput phenotyping, Hyperspectral imaging, Phenomics, Waterlogging

## Abstract

Waterlogging is a major constraint on barley productivity, yet its dynamic, multi-phase nature makes it challenging to dissect using traditional phenotyping approaches. High-throughput phenotyping (HTP) platforms address this by enabling temporal, multi-sensor imaging of large populations, but generate complex datasets that demand new analytical frameworks. Here, we imaged 230 barley accessions over 14 days of waterlogging stress and seven days of recovery using visible, chlorophyll fluorescence, and hyperspectral sensors. Explainable AI was applied to classify stress responses into early stress, late stress, and recovery phases, achieving 86% classification accuracy, and to identify the hyperspectral indices most informative for each phase. Water index (WATER1) and structure insensitive pigment index (SIPI) emerged as primary predictors of stress response. Longitudinal genome-wide association studies (GWAS), using a treatment-by-marker interaction model, identified 236 significant loci across 12 linkage disequilibrium blocks, implicating candidate genes involved in oxidative stress regulation, transcriptional control, and auxin transport. MYB transcription factors were consistently identified across all stress phases, underscoring their central role in waterlogging adaptation. To support interpretation of longitudinal GWAS results, we developed 3D-QTLVis, an interactive visualisation tool that extends Manhattan plots across time, enabling clearer identification of dynamic genomic regions underlying stress tolerance.

## Introduction

Plant phenotyping is traditionally based on labour-intensive and low-throughput measurements of traits (e.g., yield components) that are not well suited to capturing dynamic stress responses over time (Huang *et al*., 2014; Gill *et al*., 2019). High-throughput phenotyping (HTP) platforms address many of these limitations by enabling non-destructive, multi-sensor imaging of large plant populations at high temporal resolution (Araus and Cairns, 2014; Das *et al*., 2018). The use of imaging sensors enables the daily or even hourly quantification of key plant traits, including architecture, pigment status, water content, and photosynthetic performance (Andrade-Sánchez *et al*., 2014; Kharraz and Szabó, 2023). However, HTP advances come at the cost of data complexity: HTP generates massive, high-dimensional, and strongly time-dependent datasets that challenge conventional statistical analysis and visualisation pipelines.

Artificial intelligence (AI) has become an important tool for analysing the complex, high-dimensional datasets generated by modern plant phenotyping platforms. For example, supervised methods — including support vector machines, tree-based ensembles, and deep neural networks (DNNs) — have been used to classify stress states, segment plant organs, and predict quantitative traits from imaging and environmental data (Jordan and Mitchell, 2015; Suthaharan and Suthaharan, 2016; Szeliski, 2022). While AI offers strong predictive power, its utility in plant biology is dependent on interpretability. Specifically, researchers need the ability to determine which traits influence the stress prediction and how these traits link to stress-response mechanisms. This necessity has driven the adoption of Explainable AI (XAI). XAI algorithms and feature selection frameworks have gained traction in the research community to improve data interpretation. AI algorithms such as recursive feature elimination (RFE), permutation importance, and Shapley additive explanations (SHAP) help isolate informative traits, quantify their relative influence, thereby providing interpretable insights into plant stress responses (Wu *et al*., 2022; Wang *et al*., 2025).

In this study, we focus on barley-a major cereal crop with multi-uses (food, feed and malting). Barley is particularly sensitive to waterlogging, which typically occurs in regions with high rainfall or poorly drained soils (Holden *et al*., 2003; Gupta *et al*., 2022). Waterlogging limits root zone oxygen availability, severely impacting growth and yield (Lesk *et al*., 2016; Borrego-Benjumea *et al*., 2020). Although breeding for waterlogging tolerance is crucial, progress has been impaired by its complex genotype- and context-dependent nature. It is strongly influenced by external factors (soil type, stress timing, duration) and physiological traits (aerenchyma formation, root system architecture) (Luan *et al*., 2022; Ploschuk *et al*., 2023). These complexities underscore the need to discover the physiological and genetic mechanisms that confer dynamic tolerance. Barley exhibits high genetic diversity, enabling the use of several populations for association mapping (AM). Genome-Wide Association Studies (GWAS) have been the go-to method to dissect the genetic architecture of complex traits by identifying genetic variants across diverse populations that are statistically associated with a specific phenotype. Several such studies have focused on HTP in response to abiotic stress in barley (e.g., Honsdorf *et al*., 2017; Saade *et al*., 2020). In contrast, previous barley waterlogging research have focused on traditional phenotyping, thereby not capturing the crucial temporal dynamics during stress onset, maintenance, and recovery phases (Borrego-Benjumea *et al*., 2021; Luan *et al*., 2022; Manik *et al*., 2022). Recent work has established robust HTP protocols for monitoring waterlogging responses in barley (Langan *et al*., 2024), cementing its suitability for detailed characterisation of complex temporal responses and stress-induced physiological changes. The present study uses the European HerItage Barley collecTion (ExHIBiT), a genetically diverse panel of two-row spring barley representing major breeding programmes and ecogeographical backgrounds (Bernád *et al*., 2024). The ExHIBiT collection features broad morphological, physiological, and developmental variation, making it an ideal resource for exploring genotype-dependent responses to abiotic stress (Bernád *et al*., 2024).

In this study, we integrate HTP, GWAS, and XAI to dissect the dynamic waterlogging tolerance responses in barley. Using the ExHIBiT core collection in a controlled HTP experiment, we monitored plants over 14 days of waterlogging and 7 days of recovery using Red, Green, Blue (RGB), chlorophyll fluorescence, and hyperspectral imaging. From these imaging data, we derived a comprehensive suite of structural and physiological indices, including projected shoot area (PSA), chlorophyll fluorescence quantum yield (QY), and key hyperspectral indices such as Water Index 1 (WATER1) and Plant Senescence Reflectance Index (PSRI), which capture changes in water status and senescence dynamics. Next, we applied XAI to classify stress phases to identify the minimal, most reliable set of traits capturing the temporal waterlogging response. We then performed longitudinal GWAS of these key traits across multiple time points using an interaction model (Al-Tamimi *et al*., 2016). This analysis identified critical loci and candidate genes associated with specific phases (early stress, late stress, and recovery), highlighting functional roles in processes such as oxidative stress regulation, hormone signalling, and auxin transport. Finally, we address the challenge of visualisation of longitudinal GWAS results by developing an interactive tool three-dimensional genomic visualizer (3D-QTLVis), which extends conventional Manhattan plots by adding time as a third axis. 3D-QTLVis allows users to explore temporal patterns of marker effects, and examine allelic effects on phenotypes, thereby facilitating the interpretation of complex longitudinal datasets. Together, the integration of HTP, longitudinal GWAS and XAI has the potential to accelerate genomics-assisted breeding for stress tolerance.

## Methods and materials

### Plant material and genotypic data

This study used the ExHIBiT core collection, which consists of 230 spring two-row barley cultivars (Bernád *et al*., 2024). From the 50K iSelect SNPs array data (Bayer *et al*., 2017), 24,876 high-quality markers were retained by removing those with a call rate lower than 90% and a Minor Allele Frequency (MAF) greater than 5%. Marker positions were determined using the “Morex V3” reference (Mascher *et al*., 2021). Population structure analysis was conducted using STRUCTURE V.2.3.4 software (Pritchard *et al*., 2000). The assessment of LD decay and LD blocks for each chromosome were determined using the ’Ldheatmap’ R package (Shin *et al*., 2006). Further details on the collection and analysis can be found in (Bernád *et al*., 2024).

### Image-based phenotyping and experimental design

The HTP experiment was carried out in University of Picardie Jules Verne (UPJV), France using a PlantScreen™ Modular Facility (Photon System Instruments - PSI, Drásov, Czech Republic). Seeds of the ExHIBiT core collection were germinated on Whatman paper before being transferred into three litre double-pots. Drilled pots (RP 3L, Ippoland, Poland) were filled with 1.22kg of nutrient topsoil (NF U 44-551, Botanic, France) and placed in un-drilled pots (KR-3 LTR, 15cm, Kreuwel Plastics, Netherlands). Controlled conditions were assessed by environmental sensors with a day/night photo-period of 16H/8H and temperatures of 19°C/15°C with an average relative humidity maintained at 60%. Blue mats with a central hole (147mm of diameter) and plant holders (for 3L-pots, 305mm Height) were provided by PSI and used to improve image segmentation.

The imaging suite consisted of eight lanes, each one with 22 positions, with each position holding a tray containing a single pot and plant. Control and waterlogged trays were placed adjacent to each other in the growing facility to minimise spatial variation and following a split-plot design. The arrangement of control and waterlogged plants alternated within rows, and the layout of genotypes was fully randomised. All details pertaining to the number of genotypes per run, and timelines are presented in Supplementary Table S1. To account for possible spatial effects, most accessions were included in at least two different runs, with their positions rotated to different areas of the room for each run. Some level of overlap between runs was intentionally maintained to allow for cross-run comparisons.

Daily imaging was performed on all plants from one day prior to stress imposition until harvest using the PlantScreen™ Modular System (Photon System Instruments (PSI), Drásov, Czech Republic) and was conducted during the day at 1:00 pm for RGB, VNIR and SWIR images. RGB images were captured using two side-facing cameras (RGB1: 0 and 90° - GigE µEye UI-5580SE-C/M - 5 Megapixels QSXGA Camera with 1/2” CMOS Sensor) and one top-down camera (RGB2: GigE µEye UI-5580SE-C/M - 5 Megapixels QSXGA Camera with 1/2” CMOS Sensor) positioned 500 mm above each tray at a central location. The hyperspectral images acquisition was performed using two cameras in the range of visible and near-infrared spectrum (VNIR, 380 – 870 nm) and shortwave infrared spectrum (SWIR, 930 – 1670 nm). A full list of indices, their full name, description and equation can be found in Supplementary Table S2 Chlorophyll fluorescence images were captured during the night at 4:00 am to ensure full dark adaptation of the plants, using the Kinetic Chlorophyll Fluorescence Imaging Unit of the PlantScreen™ Modular System (1.4 Megapixel high-quality CCD sensor, 35 × 35 cm). The Fv/Fm protocol was employed, with an F0 measurement lasting 2 seconds, followed by an 800 ms saturating pulse set at 20% (1137 µE of intensity). The variable fluorescence (Fv) was calculated as the difference between Fm and F0.

Waterlogging stress was applied at the third leaf stage (Zadoks 13) following the protocol established by (Langan *et al*., 2024). The water level was maintained approximately 1 cm above the soil surface, corresponding to around 140% of field capacity, while control pots were kept well-watered at about 100% of field capacity. After 14 days of stress, the outer undrilled pot was removed from both control and waterlogged pots to allow excess water to drain, initiating the 7-day recovery phase, during which soil saturation was returned to 100% for all pots. The watering regimes were monitored daily at 8:30 am using the high-precision irrigation system of PSI, which is fully automated and regulates watering based on reference weight.

### Destructive phenotyping

Dry mass for three plants per watering condition and per run were measured on the final day of imaging. Samples were dried in an incubator (ULM500 Memmert) at 70°C for 48 hours and weighed using a precision balance (SBA52 Scaltec). PSA was calculated as the sum of segmented pixels from the two side angles (0° and 90°) and the top view images as previously described by (Berger *et al*., 2012). To correlate it with pixel count, a regression analysis was performed using dry weight and PSA.

### Image Segmentation

Our imaging-related datasets relied heavily on image segmentation to extract plant data due to the substantial background noise present in each image. However, not all imaging sensors required this step. Data from chlorophyll fluorescence and RGB images of the PSI PlantScreen™ system provided reliable plant masks requiring no major adjustments. In contrast, the VNIR and SWIR hyperspectral images required re-segmentation As a result, segmentation was carried out using the botanic spectrum analyser (BSA), which is a graphical user-interface (GUI) deep learning tool designed for plant image segmentation (Walsh *et al*., 2025). Therefore, the BSA was used to generate the binary plant-background masks applied to all VNIR and SWIR images in our datasets.

The goal of our segmentation process was to develop an autonomous pipeline capable of accurately segmenting plants at any growth stage. Early segmentation tests (Fig. 1) highlighted the difficulty of creating a single framework that could handle both small and large plants, especially given the temporal scale of the experiment and the 230 barley genotypes involved. This required a method adaptable to wide variation in plant size, shape, and genetics. To address this, we used the deep neural networks implemented in BSA to learn robust plant–background distinctions across diverse growth stages, and we trained and tested these models to evaluate their ability to generalise across genotypes, maintain segmentation accuracy over time, and outperform more conventional threshold- or rule-based approaches. To evaluate segmentation accuracy, a small set of reference masks was manually created with input from domain experts. Between 10 and 20 images from the RGB, VNIR, and SWIR datasets were hand-segmented using open-source image-editing tools, providing reliable benchmarks against which BSA model outputs could be compared.

**Fig. 1.**
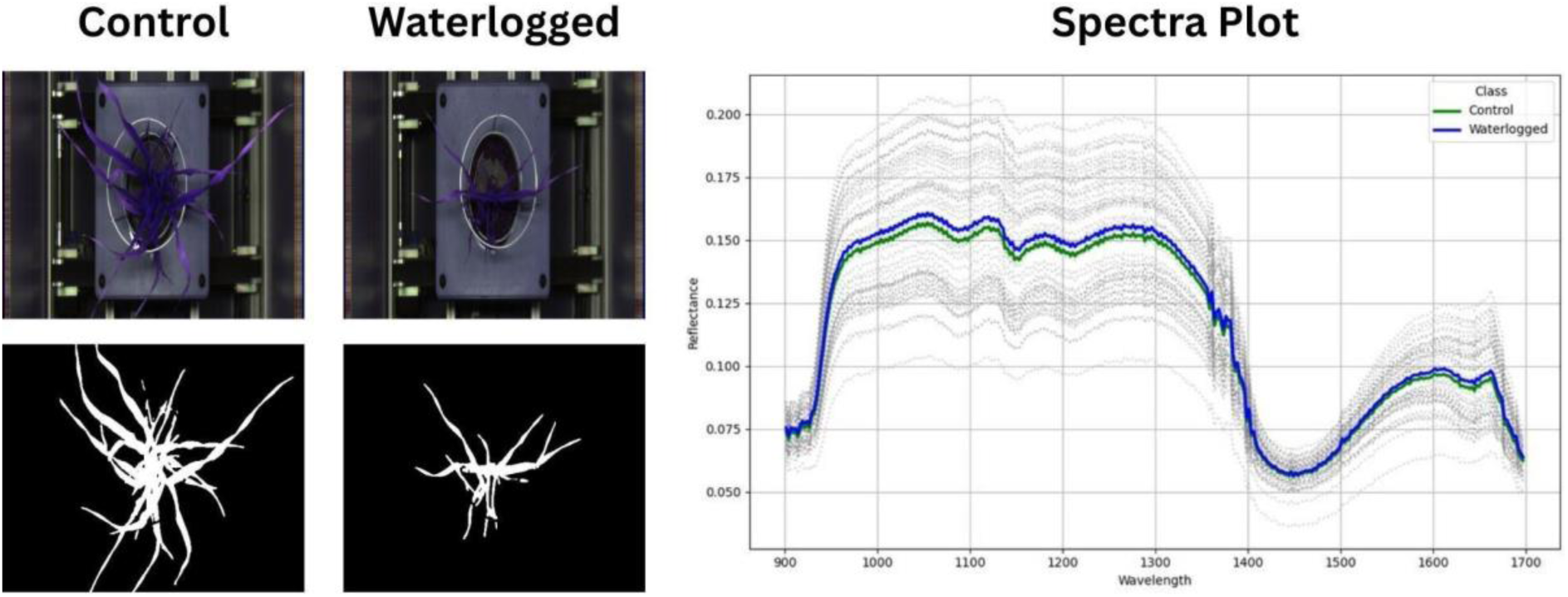
Representative segmentation outputs and mean spectral signatures for control and waterlogged barley plants. Top panels show example RGB slices of Short-Wave Infrared (SWIR) images for each treatment, with corresponding binary plant–background masks displayed below. Segmentation of SWIR hyperspectral images was performed using the Botanic Spectrum Analyser (BSA) software. The right panel presents reflectance spectra (900–1700 nm) derived from segmented hyperspectral data, where thin grey lines represent individual plant spectra and thick coloured lines indicate treatment means (n = [391] control plants; n = [391] waterlogged plants). **Alt text:** Composite figure with four panels. Left panels show SWIR hyperspectral images and corresponding binary segmentation masks for one control and one waterlogged barley plant. Right panel shows overlaid reflectance spectra from 900 to 1700 nanometres, with individual plant spectra in grey and treatment means as bold lines, comparing control and waterlogged conditions.

### Data Analysis

A linear mixed model was fitted for all traits, except for PSA, to combine the six independent runs. Model consisted of following terms: y(ijk) = genotype(i) + treatment(j) + genotype * treatment(ij) + run(k) + error, with the terms in bold being the random effects, while all others were fixed, adopted from (Honsdorf *et al*., 2017). In the case of the PSA data, the runs were combined by taking the average across all the runs. All trait values were smoothed by fitting a cubic smoothing spline to the data (Al-Tamimi *et al*., 2016). For PSA, these smoothed values were used for all subsequent analysis while for other traits these values were only used for visualisation purposes. Relative Growth Rate (RGR) was calculated using Equation 1 across two-time intervals, day-to-day and using the time intervals identified by the unsupervised clustering. The Pearson correlation coefficient was calculated using the R package *Hmisc* (Harrell, 2023) within the R statistical computing environment R v3.6.0 (R Core Team, 2021). The resulting correlation matrix was then visualised with the corrplot package in R (Wei and Simko, 2021). Heritability was estimated following the approach described by (Cullis *et al*., 2006). Statistical significance was assessed using either ANOVA or the Kruskal-Wallis test (Kruskal and Wallis, 1952). Data visualisation was performed with the R package ggplot2 (Wickham *et al*., 2016).

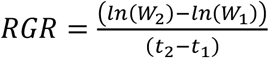

### AI for key trait identification and time-responses

To facilitate the interpretation of the plants’ dynamic responses to waterlogging, we applied unsupervised and supervised AI algorithms to our 21-day time series datasets (RGB, VNIR, SWIR). Our goal was to determine whether temporal patterns could be grouped into biologically meaningful phases that would simplify AI and GWAS analysis.

First, we used unsupervised clustering to determine whether the 21-day time series exhibited inherent temporal structure. We evaluated several clustering algorithms (k-mean, hierarchical clustering, and DBSCAN) and visualised their separability using dimensionality-reduction methods (Principal Component Analysis (PCA) and t-Distributed Stochastic Neighbour Embedding (t-SNE). Cluster evaluation metrics, including the elbow method and the silhouette index, were computed to estimate the optimal number of clusters. These analyses were performed on both the full phenotypic dataset and on a derived “tolerance dataset” that highlighted genotype-specific differences between waterlogged and control conditions.

Second, because unsupervised clustering alone could not resolve all stress-response patterns, we complemented this step with supervised learning. The three temporal phases identified through clustering were used as class labels for training a suite of machine learning (ML) and deep learning (DL) algorithms. The primary purpose of this classification analysis was not to define the phases, but to assess their discriminability and identify the phenotypic traits most informative for distinguishing them.

Finally, to identify the traits most informative for distinguishing waterlogging stress, we applied a set of feature-importance algorithms. RFE was used to remove redundant variables and retain a compact set of predictors, followed by permutation importance to quantify each trait’s contribution to classification accuracy. To further enhance interpretability, SHAP values were computed for selected algorithms, providing class-level insights into how individual traits influenced predictions. Together, these approaches allowed us to determine which traits contributed most strongly to the detection and characterisation of waterlogging stress.

### GWAS and dynamic results’ visualisation with 3D-QTLVis

To identify loci associated with waterlogging response, GWAS was performed on the ExHIBiT core collection using the imaging traits identified during the Feature importance analysis. Two GWAS models were utilised: Bayesian-information and Linkage-disequilibrium Iteratively Nested Keyway (BLINK) model (Xia *et al*., 2019), implemented through the Genome Association and Prediction Integrated Tool package (GAPIT) package (Wang and Zhang, 2021) and Interaction model (Al-Tamimi *et al*., 2016). The first three principal components (PCs) were included alongside a kinship matrix for genetic relatedness correction in both models. Marker-trait association significance was determined based on the False discovery rate (FDR) threshold of >0.05 and a Bonferroni-adjusted threshold (**α** = 0.05). Two versions of the Interaction model were developed using standard R (R Core Team, 2021) as a user-friendly tool for the community, R scripts can be found in https://github.com/Walshj73/3D-QTLVis. The model ‘Int+Q’ included only population structure whereas ‘Int+Q+K’ included both population structure and kinship matrix. Both models were applied in case the inclusion of both population structure and kinship caused overfitting of the model.

The genomic inflation factor (λ) was calculated as the ratio of the median observed chi-square statistic to the expected median under the null hypothesis. This analysis, alongside Quantile-Quantile (QQ) plots, utilised the qqPlot() function from the “car” package in R (Fox and Weisberg, 2019) and was subsequently employed to evaluate the optimal GWAS model.

To facilitate the interpretation of longitudinal GWAS results, we developed a new user-friendly visualisation tool-3D-QTLVis-that extends Manhattan plots throughout time. Our tool was implemented using the Shiny package (Chang *et al*., 2025) within the R statistical computing environment (version 3.6.0) (R Core Team, 2021). Graphs’ visualisations were developed using the Plotly package (Sievert *et al*., 2021). Full R script can be found at https://github.com/Walshj73/3D-QTLVis. 3D-QTLVis includes three interactive graphical representations to visualise GWAS data. The first visualisation is a 3D Manhattan plot, which displays the Logarithm of Odds (LOD) score against chromosome location, with a third axis added to represent time. The second visualisation is a line graph, which shows the variation in LOD score for a specific Single Nucleotide Polymorphism (SNP) across the time points in the study. The third visualisation is a boxplot, which illustrates the phenotypic variation of the selected SNP across its alleles. Graphs 2 and 3 are dynamically generated and appear when a user clicks on a specific SNP of interest within the main 3D Manhattan plot. The data required for these visualisations include the GWAS results (specifically the p-values from the GWAS analysis) as well as phenotypic data.

### Candidate gene selection

Candidate gene selection prioritized loci exhibiting high heritability, high effect sizes, and traits linked to significant Linkage Disequilibrium (LD) blocks identified at multiple days and in multiple traits. Genes located within these significant LD blocks were annotated using EnsemblPlants (https://plants.ensembl.org). Furthermore, gene function was acquired from UniProt (https://www.uniprot.org/).

## Results

### Explainable AI improves interpretation of hyperspectral data for dynamic stress responses

We imaged the ExHIBiT core collection, consisting of 230 accessions, using a HTP platform over a 21-day period, which included 14 days of waterlogging and 7 days of recovery. The experiment was conducted in six separate runs, with some level of accession overlap between each run. The data collected was subsequently averaged for each accession. The experiment was carried out in six separate runs with shared accessions between all runs, the values then averaged per accessions. Four imaging sensors, RGB, Chlorophyll fluorescence, VNIR and SWIR were used to collect data. A total of 11 indices were derived, including PSA and RGR using the RGB sensors, QY from chlorophyll fluorescence sensors, Normalised Difference Vegetation Index (NDVI), Normalised Difference Vegetation Index 2 (NDVI2), Photochemical Reflectance Index (PRI), Structure Insensitive Pigment Index (SIPI), PSRI, Modified Chlorophyll Absorption in Reflectance Index 1 (MCARI1), and Optimised Soil-Adjusted Vegetation Index (OSAVI) using the VNIR camera, and WATER1 using the SWIR sensor (Supplementary Table S2). We found that waterlogging stress significantly impacted all indices, with a dynamic variation observed across time points and between treatments. Except for WATER1, all traits displayed a reduction because of waterlogging, while WATER1 showed an initial decline after day 7 before increasing compared to the control (Table 1).

**Table 1.**
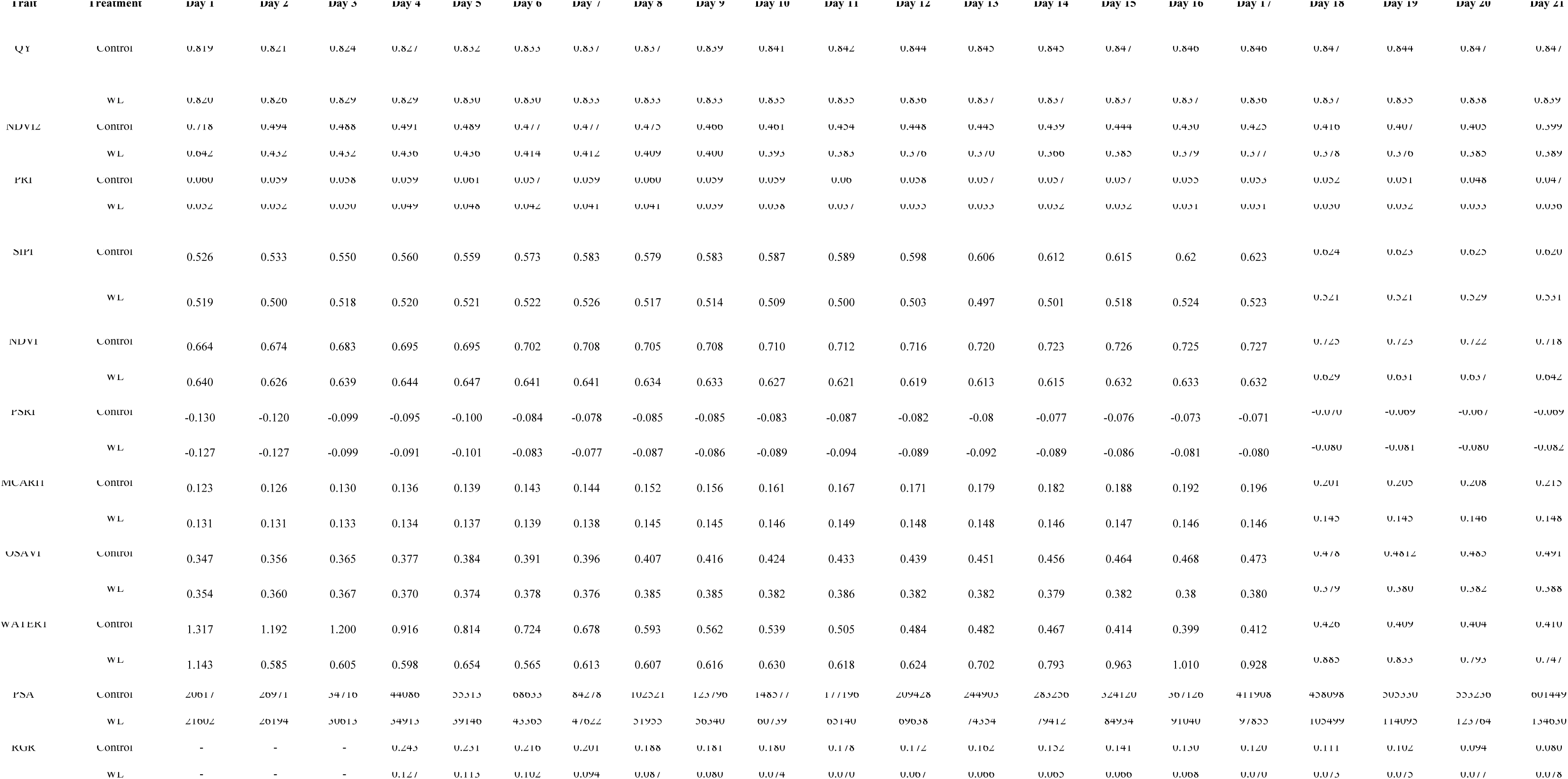
Mean values of eleven imaging barley phenotypic traits under control and waterlogging stress (WL) conditions over a period of 14 days of waterlogging and 7 days of recovery. Traits are as follows: quantum yield (QY), Normalised Difference Vegetation Index 2 (NDVI2), Photochemical Reflectance Index (PRI), Structure Insensitive Pigment Index (SIPI), Normalised Difference Vegetation Index (NDVI), Plant Senescence Reflectance Index (PSRI), Modified Chlorophyll Absorption in Reflectance Index 1 (MCARI1), Optimised Soil-Adjusted Vegetation Index (OSAVI), Water Index 1 (WATER1), Projected Shoot Area (PSA), and Relative Growth Rate (RGR). Values represent means across [n = 230] accessions per treatment [averaged across biological replicates from six independent runs].

The RGR captured day-to-day changes in growth rates (Supplementary Fig. S1). Early measurements were excluded due to segmentation challenges in small plants, which occasionally resulted in negative RGR values (Table 1). In the control group, RGR was initially high, exhibiting a steady decline over time. In waterlogged (WL) plants, RGR also decreased initially, but began to recover at the start of the recovery phase. By the end of the experiment, WL plants demonstrated a resurgence in growth, with RGR values approaching those of the control group. Because our GWAS analysis did not show significant associations with RGR, we have opted to omit it from further exploration in our study, which also applies to QY. To facilitate longitudinal data interpretation, we used AI algorithms to identify an appropriate temporal structure within the stress response. Unsupervised clustering of the 21-day time series revealed that the daily measurements did not form distinct groups but instead aggregated into a smaller number of recurring temporal patterns. Initial k-means analysis suggested 2–4 clusters, and PCA and t-SNE visualizations indicated substantial overlap when using the full phenotypic dataset. In contrast, clustering analysis yielded a clearer structure, with k = 3 producing the highest silhouette and the most coherent grouping. Based on this outcome, we partitioned our time series dataset into three temporal phases: early stress (1-7 days), late stress (8-14 days), and recovery (15-21 days).

Subsequently, we leveraged explainable AI methods for feature selection, determining the most predictive traits from the 11 examined HTP indices. Using these three phases as class labels, we evaluated the performance of multiple AI algorithms. Most algorithms achieved initial weighted F1-scores above 70%, with logistic regression being the only method below this threshold. Three-based and boosting algorithms (e.g., Extreme Gradient Boosting (XGBoost), Categorical Boosting (CatBoost) showed strong baseline performance (>70%) and only minor improvements following hyperparameter optimisation (Table 2).

**Table 2.** Weighted F1-scores for various artificial intelligence (AI) algorithms applied to barley dataset containing extracted phenotypic traits, ranked from highest to lowest performance. The feed-forward neural network (FNN) algorithm achieved the highest weighted F1-score, indicating superior performance in balancing precision and recall across stress phases.

| Algorithm | Weighted F1-score (%) |
| --- | --- |
| Feed-forward Neural Network | 86 |
| Support Vector Machine | 83 |
| CatBoost | 78 |
| XGBoost | 78 |
| Decision Tree | 76 |
| K-Nearest Neighbour | 75 |
| Random Forest | 72 |
| Logistic Regression | 62 |

The feed-forward neural network (FNN) achieved the highest performance, with an initial weighted F1-score of 84% increasing to 86% after increasing the network depth and filter size. Support vector machine (SVM) and k-nearest neighbours (KNN) exhibited the largest gains from optimisation, with SVM improving from 40% to 83%. Confusion matrices across models consistently showed misclassification of late stress, primarily as recovery and, to a lesser extent, early stress. To determine which traits contributed to these classification outcomes, we next evaluated feature importance across algorithms. Feature-importance analyses indicate that PSA was the most influential trait in all models, contributing approximately 40–50% of predictive accuracy. Removal of PSA resulted in a substantial performance decrease across algorithms. Permutation-based ranking confirmed PSA as the top predictor, followed by OSAVI and MCARI1. SHAP analyses (Fig. 2) for the best-performing models (SVM and FNN) further identified PSA and SIPI as the main contributors for waterlogging responses across stress phases, with WATER1 showing moderate importance, particularly for early stress.

**Fig. 2.**
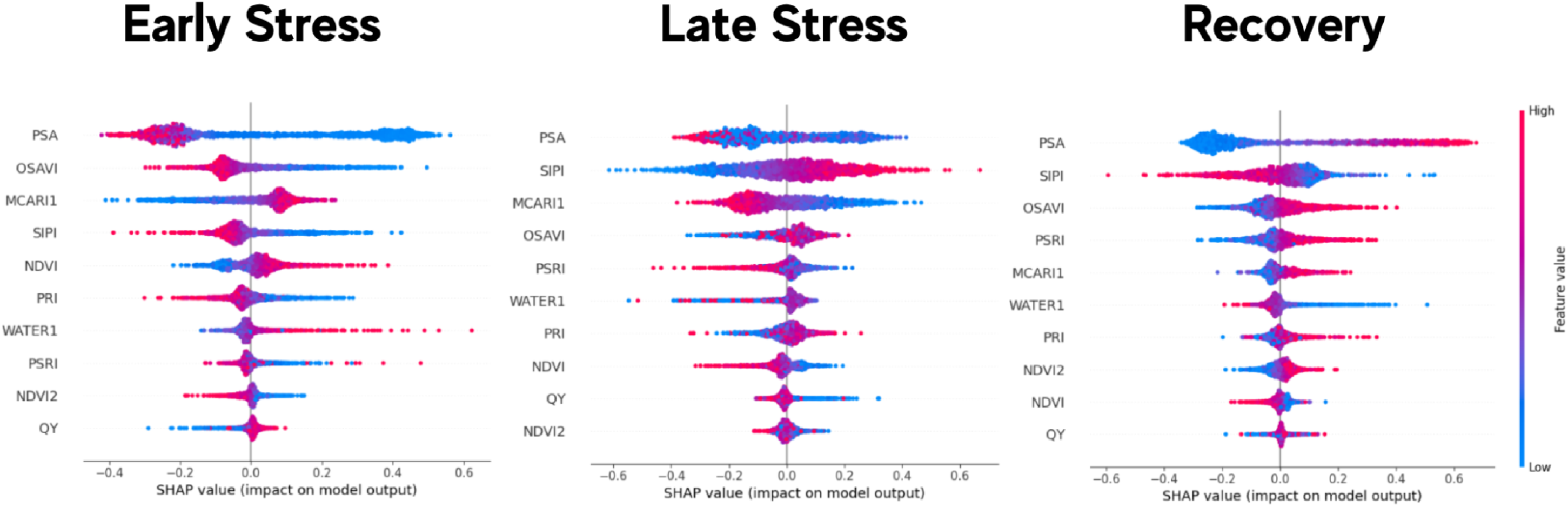
SHAP (Shapley Additive Explanations) summary plots showing the contribution of vegetation indices to model predictions during three waterlogging stress phases: early stress (1–7 days), late stress (8–14 days), and recovery (15–21 days). The x-axis represents SHAP values (feature contributions to the model output). The vertical line at the centre of each panel represents a SHAP value of zero (no effect on the model prediction). Points to the right of zero indicate a positive contribution to the prediction, whereas points to the left indicate a negative contribution. The colour gradient (blue to red) represents feature magnitude, with blue indicating lower feature values and red indicating higher feature values (n = [2912] observations per phase, derived from [230] accessions across biological replicates). **Alt text:** Three horizontal beeswarm plots side by side, each representing one stress phase (early stress, late stress, and recovery). Each plot ranks vegetation indices vertically by importance and displays individual SHAP values as coloured dots along a horizontal axis, with colour indicating feature value from low (blue) to high (red).

Next, we confirmed that PSA values accurately reflect plant growth by comparing them with destructive phenotyping data. The strong correlation (R2 = 0.92) observed across all runs confirms the reliability of our imaging and analysis workflow (Supplementary Fig. S2). The PSA showed significant differences between control and WL plants from day 1 until day 21. Under control conditions, PSA values increased steadily, consistent with healthy growth. In contrast, WL plants exhibited significantly lower PSA values by day 21, indicating marked growth inhibition (Fig. 3).

**Fig. 3.**
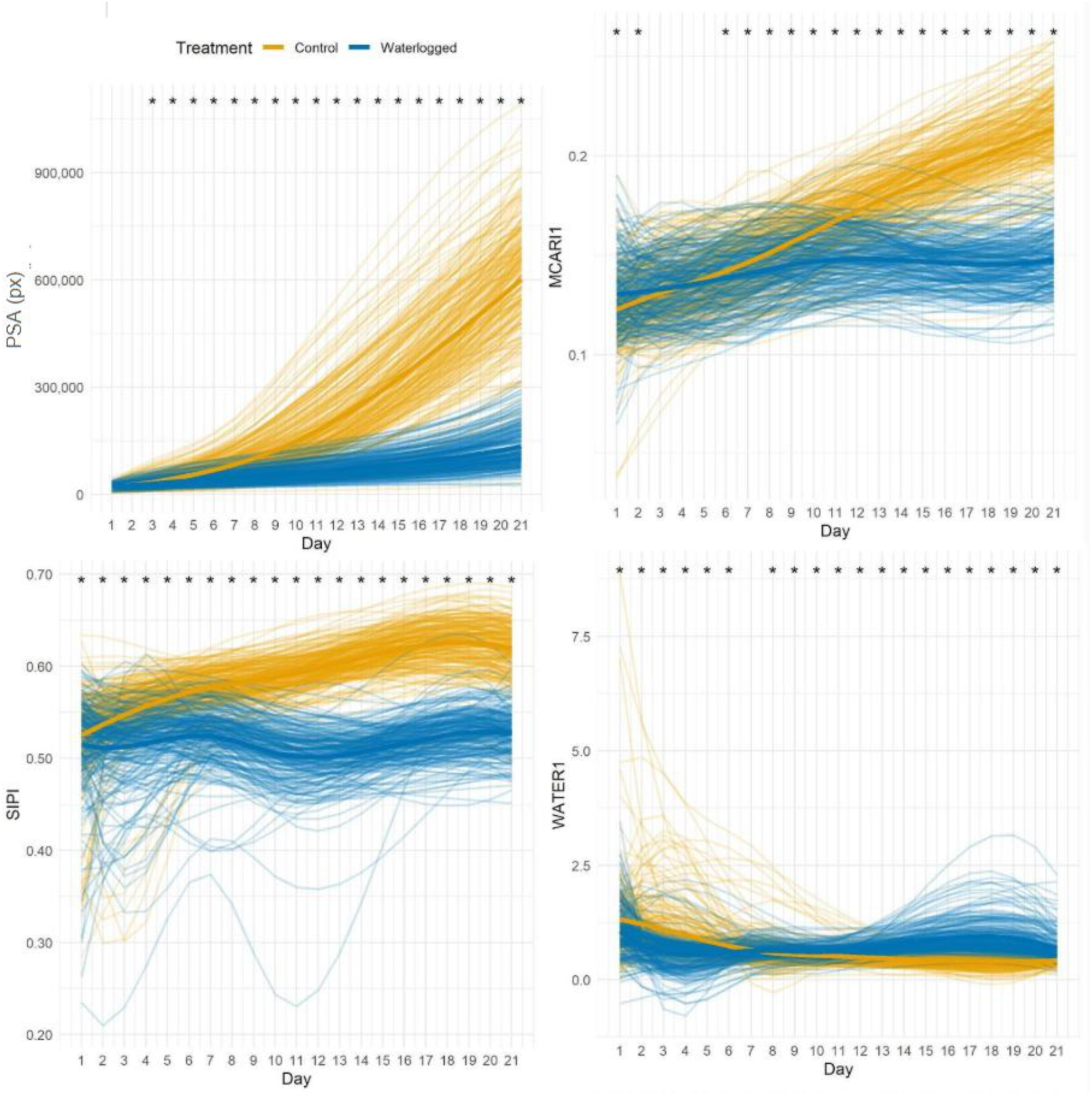
Polynomial curves of imaging traits over 14 days of waterlogging followed by 7 days of recovery in 230 barley accessions. Yellow lines represent the control treatment, while blue lines represent the waterlogged treatment. Each thin line corresponds to an individual accession averaged across biological replicates from separate runs, and the bold lines depict the mean values for each treatment (n = [391] control; n = [391] waterlogged). Black stars indicate statistically significant differences between control and waterlogged treatments on specific days, determined using [ANOVA/Kruskal-Wallis test] (two-sided, P < 0.05). (A) Projected Shoot Area (PSA), (B) Water Index 1 (WATER1), (C) Structure Insensitive Pigment Index (SIPI), (D) Modified Chlorophyll Absorption in Reflectance Index 1 (MCARI1). **Alt text:** Four line graphs arranged in a two-by-two grid, each showing a different vegetation index over 21 days. Each panel displays individual accession trajectories as thin lines and treatment means as bold lines, with yellow for control and blue for waterlogged. Black stars mark days with statistically significant treatment differences.

Our results demonstrate the efficacy of hyperspectral sensors and their derived vegetation indices in characterising stress responses. We observed substantial correlation among all evaluated indices. The highest correlations were found between MCARI1 & OSAVI (r = 0.97), SIPI & NDVI (r = 0.93) and MCARI1 & PSA (r = 0.81). The full correlation matrix can be seen in Supplementary Fig. S3 WATER1 has low correlation with the other traits due to being derived from a different hyperspectral camera. By using explainable AI, we confirmed that three specific hyperspectral indices were the most informative traits characterising the dynamic responses of barley to waterlogging. The first index, MCARI1 shows key differences, with significant variation detected between days 1-2 and again from Days 6 to 21 (Fig. 3). Initially, control values were lower than those under waterlogging, but by day 4, control values surpassed WL and remained higher for the remainder of the experiment. The second index, SIPI, demonstrates significant differences between control and WL treatments across all time points from day 1 to day 21 (Fig. 3). Control plants consistently displayed higher SIPI values than WL plants, with the difference gradually increasing over the span of the experiment, suggesting a cumulative effect of waterlogging stress on this index. Following the onset of recovery, the gap in SIPI values between control and WL plants narrowed slightly, indicating a partial recovery response. The final key index, WATER1, displays significant differences between control and WL treatments across days 1-6 and 8-21 (Fig. 3). Initially, control plants exhibited higher WATER1 values than WL plants. However, as the experiment progressed, WATER1 values in WL plants increased steadily, eventually surpassing those of the control by mid-experiment. Notably, following the start of the recovery phase, WATER1 values in WL plants began to decline. Response to waterlogging stress and general trends for the rest of the imaging traits can be seen in Supplementary Fig. S4-S8.

To evaluate the proportion of phenotypic variance that is due to genetic differences between individuals, we examined heritability values across traits and treatments (Supplementary Table S3). Our results show that PSA exhibited moderate heritability under control conditions (0.5174), which increased notably under waterlogged conditions (0.7219). SIPI showed moderate heritability, with slight reductions under waterlogging (0.4525 control vs. 0.3848 WL). NDVI showed consistent heritability between conditions (0.4118 control vs. 0.4215 WL). MCARI1 displayed moderate heritability both under control (0.5078) and under WL conditions (0.6281). Finally, WATER1 showed low heritability values under both control (0.1869) and WL (0.1530) conditions.

### GWAS Interaction model improves the dissection of complex stress phenotypes

To examine the dynamic quantitative traits related to waterlogging stress in barley, we conducted GWAS throughout time. Our results identified a total of 4,983 markers associated with 11 traits derived from four HTP imaging sensors. After a series of filtering steps, LD blocks associated with a single trait at a single time point were removed, as were duplicates identified across interaction models. Additional exclusions included LD blocks linked only to control treatments and blocks with fewer than two or non-continuous time points or traits. This rigorous filtering process narrowed the dataset to 236 SNPs distributed across 12 LD blocks. Notably, only the Interaction model was able to identify Quantitative Trait Loci (QTLs) over more than two consecutive days, which resulted in the majority of significant SNPs being identified through this model. A brief overview of results can be found in Table 3 and a detailed list available in Supplementary Table S4.

**Table 3.** Summary of genome-wide association studies (GWAS) results listing the linkage disequilibrium (LD) blocks with significant marker-trait associations, number of QTLs within and traits of association divided into three experimental phases: early stress, late stress and recovery. In brackets are the days for which there were significant QTLs.

| LD block | Number of significant QTLs | Early Stress (1-7 days) | Late Stress (8-14 days) | Recovery (15-21 days) |
| --- | --- | --- | --- | --- |
| H1-27 | 6 |  |  | WATER1(16,18-21) |
| H2-157 | 7 | WATER1(2-5) |  |  |
| H2-333 | 4 |  |  | PSRI(18-21) |
| H3-396 | 5 |  |  | PSRI(18-21) |
| H5-114 | 11 | WATER1(2-5) |  |  |
| H5-115 | 74 | WATER1(2-5) |  |  |
| H5-128 | 33 | WATER1(2-5) |  |  |
| H5-507 | 4 | WATER1(2-5) |  |  |
| H5-509 | 4 | WATER1(2-5) |  |  |
| H5-515 | 51 | WATER1(2-5) |  |  |
| H5-91 | 29 | WATER1(2-5) |  |  |
| H6-241 | 8 |  |  | PSRI(16-18,21) |

We used XAI to identify the key hyperspectral indices that effectively discriminate waterlogging stress. Interestingly, we found that both XAI and our GWAS Interaction model pinpointed WATER1 as having a major role in early stress responses. We found significant SNP associations with WATER1 primarily during early stress phases (days 2-5) but also in later stress phases (days 16, 18-21). Notably, in LD block H1-27, the SNP-JHI-Hv50k-2016-5784 was associated with WATER1 during the later stages. In the H2-157 block, SNP - JHI-Hv50k-2016-80221 and JHI-Hv50k-2016-80251 were significantly associated with WATER1 during early stress periods. The H5 region emerged as a hotspot for WATER1, with multiple loci (H5-91, H5-115, H5-128, H5-507, H5-509, and H5-515) showing strong associations during days 2-5. For instance, significant SNP such as JHI-Hv50k-2016-361178, JHI-Hv50k-2016-299910, and JHI-Hv50k-2016-362931 were linked to this early stress phase.

Additionally, our GWAS Interaction model highlighted PSRI as a hyperspectral index associated with the waterlogging recovery phase. PSRI was predominantly associated with SNP during the recovery phase of stress. In the H6-241 block, SNP - JHI-Hv50k-2016-422491 and JHI-Hv50k-2016-422681 were significantly associated with PSRI during days 16, 17, 18, and 21. Similarly, the H2-333 region showed a significant association between SNP JHI-Hv50k-2016-128479 and PSRI during days 18-21. Another significant association with PSRI was observed in the H3-396 region, where the SNP - JHI-Hv50k-2016-223621 exhibited links to recovery-phase traits at days 18-21.

### 3D-QTLVis: a three-dimensional visualiser for longitudinal GWAS results

Recognising the inherent complexity and interpretation challenges associated with longitudinal GWAS results, we sought to develop a new community resource designed to simplify data visualisation and streamline interpretation to uncover hidden dynamic trends in genotype-phenotype associations. Our new tool-3D-QTLVis, is an interactive Shiny app that includes three visualisation windows to facilitate GWAS data exploration throughout time. The first visualisation window is a 3D Manhattan plot, which displays genomic position (in base pairs), time (in this case days), and LOD score to show changes throughout the experiment (Fig. 4). This extension of the traditional Manhattan plot enables users to investigate how the association between genetic variants and traits evolves across different time points throughout their experiment. These windows can be zoomed in and rotated to examine the data from different angles, and it includes two planes of significance (Suggestive threshold and the Bonferroni correction). Visualisation windows 2 and 3 are interactively generated from the first window, becoming visible when a user selects a specific SNP of interest from the primary 3D Manhattan plot. The visualisation is a line graph, which shows the variation in LOD score for a specific SNP (interactively selected by the user) across the time points in the study (Fig. 4), allowing for a more detailed analysis of the temporal dynamics of the SNP association. The third visualisation window is a boxplot, which illustrates the phenotypic variation between the two alleles of the selected SNP marker (Fig. 4). This boxplot enables comparisons between treatment groups, demonstrating how genetic variation is translated into phenotypic differences.

**Fig. 4.**
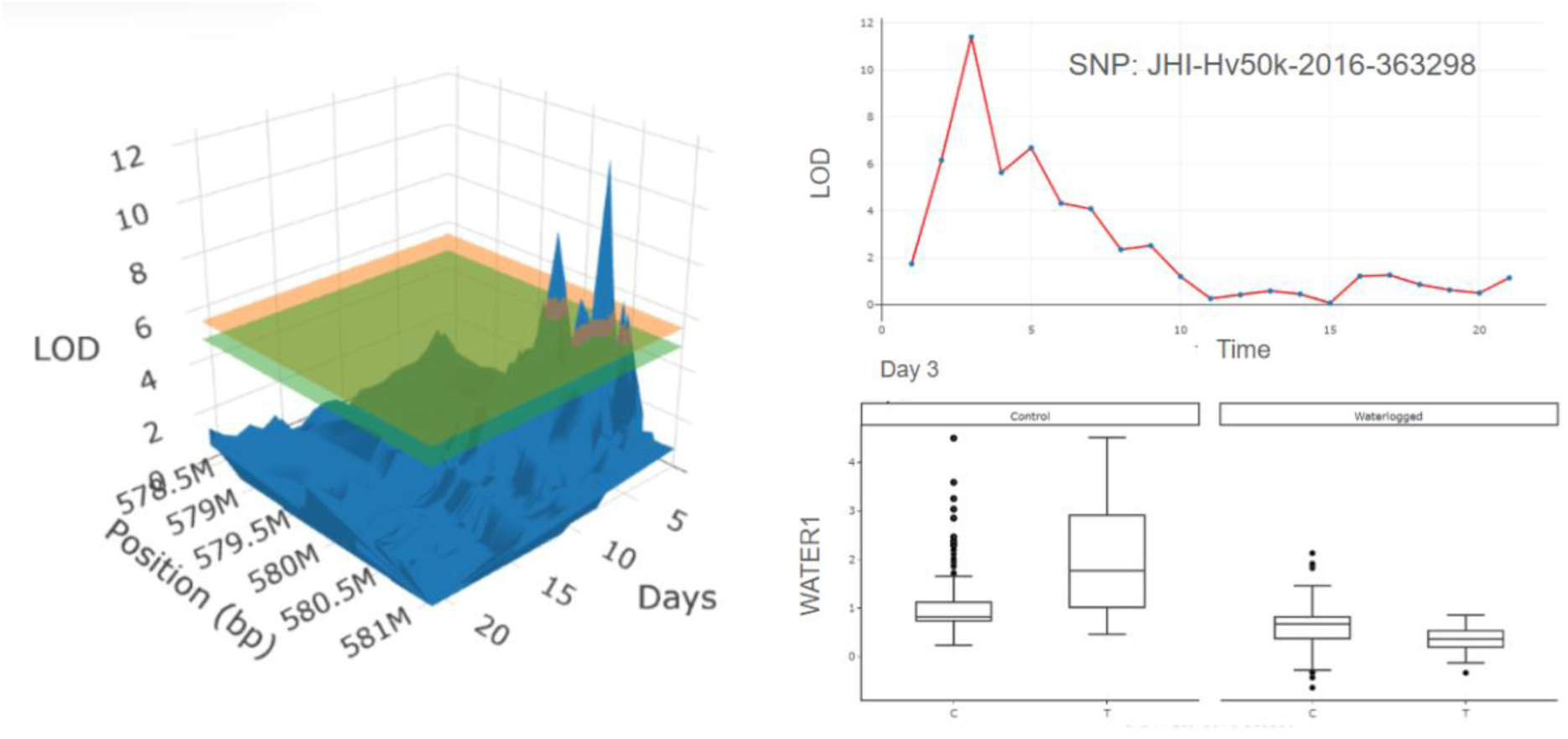
3D-QTLVis visualisation of LD block H5-515 located in barley chromosome 5, examining the genetic architecture of the spectral index WATER1. (A) 3D Manhattan plot displaying genomic position (in bp), time (in days), and logarithm of odds (LOD) score across a longitudinal GWAS analysis. The lower horizontal plane (green) indicates the suggestive significance threshold and the upper horizontal plane (orange) indicates the Bonferroni-corrected significance threshold. (B) Line graph showing the variation in LOD score for a selected SNP across the 21 time points of the study. (C) Boxplot illustrating the phenotypic variation of the selected SNP across alleles and treatments in 230 barley accessions under controlled and waterlogged conditions. **Alt text:** Three-panel figure. Panel A shows a three-dimensional Manhattan plot with genomic position, time, and LOD score as axes, with two horizontal significance threshold planes. Panel B shows a line graph of LOD score over 21 days for a single SNP. Panel C shows boxplots comparing WATER1 values across two allele classes under control and waterlogged treatments.

To illustrate the functionality of 3D-QTLVis, we used our GWAS results by highlighting one of the LD blocks of interest derived from our analysis as a concrete example. In this specific case, we examined LD block H5-515 (WATER1, Days: 2-5), where the 3D GWAS plot highlights two significant SNPs — JHI-Hv50k-2016-363298 & JHI-Hv50k-2016-362931 — that stand out and warrant closer investigation. When examining one of them in closer detail, JHI-Hv50k-2016-363298, the line graph shows a low LOD score on day 1, which increases to the highest point on day 3 and then continuously declines over time. The boxplot reveals that on day 3, WATER1 has higher values for both C and T alleles under control conditions, with values falling during waterlogging. However, the decline is more pronounced for the T allele compared to C, even though the T allele shows a higher value under control conditions but a lower value under waterlogged conditions.

### Hyperspectral indices unlock the genetic dissection of candidate genes contributing to waterlogging stress

Waterlogging tolerance in barley requires precise regulation of physiological and molecular pathways to manage stress responses. Using our GWAS Interaction model, we identified several loci with a range of genes specifically associated with stress signalling, oxidative stress defence, transcriptional regulation, and cellular transport, many of which are implicated in adaptive responses under hypoxic conditions induced by waterlogging. In total, our analysis identified 236 candidate genes across 14 LD blocks, distributed across the following LD blocks: 6 candidate genes in H1-27, 2 in H2-157, 33 in H2-333, 88 in H3-396, 21 in H5-144, 1 in H5-507, 5 in H5-509, 67 in H5-515, 78 in H5-91 and 9 in H6-241. To narrow down our search in dynamic responses to waterlogging, we focused on candidate genes with functional annotations linked to waterlogging stress, highlighting 28 candidate genes (Table S5).

The candidate genes identified in our study belong to several GO annotated categories that are known to be involved in stress responses. For example, we found auxin-related genes, such as the auxin efflux carrier component (HORVU.MOREX.r3.3HG0 327810) that is associated with PSRI. This gene is known to have a critical role in root architecture modulation, enabling improved oxygen access during waterlogging (Geldhof *et al*., 2022). We also found candidate genes associated with oxidative stress management, which are represented by glutaredoxin domain-containing proteins (HORVU.MOREX.r3.5HG0429380, HORVU.MOREX.r3.5HG0429400 and HORVU.MOREX.r3. 5HG0534650, both being associated with WATER1) and the glutathione S-transferase (HORVU. MOREX.r3.2HG0202910 associated with PSRI). According to GO terms, these genes are known to mitigate Reactive Oxygen Species (ROS) accumulation, protecting cellular structures and enzymes under hypoxia-induced stress (Mhamdi and Van Breusegem, 2018; Lana *et al*., 2024). As expected, we also found several transcription factors within our pool of candidate genes prominently associated with waterlogging stress. The identified transcription factors, include MYB transcription factors (HORVU.MOREX.r3.5HG0 428450 associated with WATER1), Myb/SANT-like domain-containing proteins (HORVU.MORE X.r3.3HG0328110 linked to PSRI), and NAC domain-containing proteins (HORVU.MOREX.r3.5 HG0533860 to 5HG0533960 associated with WATER1). MYB transcription factors are involved in secondary metabolism and cell wall reinforcement, while NAC proteins regulate processes such as programmed cell death and stress-induced morphogenesis (Mhamdi and Van Breusegem, 2018; Lana *et al*., 2024). Furthermore, our analysis revealed candidate genes involved in several key known functions contributing to waterlogging stress responses, namely peroxidase (HORVU.MOREX.r3.5HG0428280 associated with WATER1), which contributes to lignification and cell wall fortification (Mhamdi and Van Breusegem, 2018; Lana *et al*., 2024). Our study also detected protein kinase domain-containing proteins across multiple loci, which are involved in signal transduction mechanisms, including HORVU.MOREX.r3.1HG0002 680 and 2HG0202810. These proteins are involved in phosphorylation cascades that activate downstream stress-response pathways (Song *et al*., 2025). Additionally, the sucrose transporter (HORVU.MOREX .r3.2HG0203150 linked to PSRI) and WAT1-related protein (HORVU.MOREX .r3.2HG0203240 linked to PSRI) are implicated in carbohydrate partitioning and cell wall biosynthesis, both crucial for energy supply and cell wall restructuring under hypoxic stress (Shigeto *et al*., 2013; Lauschke *et al*., 2025). Collectively, the integration of hyperspectral indices in the GWAS interaction model revealed candidate genes distributed across chromosomes 1H, 2H, 3H, and 5H, representing diverse functional GO categories contributing to waterlogging adaptation in barley.

## Discussion

Breeding for waterlogging tolerance is particularly challenging due to its complex polygenic and dynamic temporal nature, which involves multiple tolerance mechanisms. Traditional phenotyping methods often fall short in addressing these needs, as their destructive nature limits their utility for monitoring temporal changes. High-throughput phenotyping using modern imaging technologies aims to bridge this gap. This study combines HTP and XAI to investigate waterlogging tolerance in barley. This integrative strategy, coupled with association mapping, ultimately led to the identification and understanding of genes associated with waterlogging stress over time.

### AI facilitates dynamic stress responses interpretation

AI has become a powerful tool for interpreting complex phenotypic datasets, particularly those that involve dynamic physiological responses (Cembrowska-Lech *et al*., 2023). Recent work emphasises that although AI models can successfully extract hidden patterns from high-dimensional phenomic data, their performance is often constrained by the inherent heterogeneity and noise that characterise plant stress responses (Koh *et al*., 2024). This is especially evident in imaging-based datasets, where factors such as illumination variability, sensor artefacts, and subtle early-stage physiological changes introduce inconsistencies that can inflate error rates when standard accuracy metrics appear favourable (Islam *et al*., 2024).

Our study used XAI to classify barley responses to waterlogging, leveraging algorithms such as KNN and SVM. We achieved robust classification accuracy across most of the algorithms (seven out of eight), providing valuable insights into how AI can be used for interpreting complex abiotic stress responses despite the known challenges inherent in applying AI in HTP. Clear signs of overfitting were visible in both tree-based algorithms and KNN, which achieved high training performance but a noticeably reduced validation accuracy. Similar patterns have been documented in other stress classification studies. For example, (Naik *et al*., 2017) found that traditional classifiers frequently overfit when canopy-level colour traits or spectral indices are noisy or highly correlated. To improve the overfitting of the algorithms, we performed regularisation and parameter tuning. These modifications were found to alleviate this overfitting behaviour but did not fully resolve it, supporting that AI algorithms have sensitivity to noise generated by HTP indices. Previous research indicates that neural networks often surpass classical ML algorithms in performance because they learn the hierarchical representations rather than relying on pre-defined features, thereby offering greater resistance to overfitting even under extreme training regimes (Gill *et al*., 2022). Our study showed that the FNN generalised well throughout training and only showed signs of overfitting under extreme training conditions, corroborating with the results from (Gill *et al*., 2022). To further improve the FNN performance, we introduced early stopping, checkpointing and dropout to stabilise the model, reflecting widely adopted strategies for improving generalisation in deep neural networks (Mostafa *et al*., 2023).

After investigating the differing behaviours of the algorithms, we sought to understand which phenotypic traits were most informative for distinguishing our waterlogging phases, i.e., early, late and recovery. Across all algorithms tested, PSA emerged as the most influential contributor across all algorithms and in all waterlogging phases, reinforcing its known sensitivity to abiotic stress. In cowpea, Yu (2024) showed that drought-stress caused significant reductions in PSA as early as four days after stress onset. Taken together, these findings demonstrate that PSA reliably reflects early disruptions in shoot and canopy development under a range of abiotic stress, which explains why it served as the dominant trait (feature) for identifying each waterlogging stress phase in our algorithms. We also observed large phenotypic variations across all traits, underscoring the diversity in stress responses within the population. Waterlogged plants exhibited marked reductions in growth dynamics compared with controls, with significant decreases detected in vegetation indices, including PRI, MCARI1, NDVI2, NDVI, OSAVI, and SIPI as well as a reduction in QY. Our results are in good agreement with previous studies showing that a decline in vegetation indices reflects compromised plant health under abiotic stress, for example (Ihuoma and Madramootoo, 2019), identified OSAVI, water index, and NDVI as the most sensitive indices for detecting drought stress in tomatoes. Jiang et al. (2013) found that SIPI and PRI were effective in identifying waterlogging stress in maize and beetroot. In addition, Römer *et al*. (2012) found that PRI could detect differences in nutrient treatments, while NDVI successfully identified drought stress in barley.

Notably, we observed significant differences in NDVI, NDVI2, PRI, SIPI, and WATER1 as early as day 1, highlighting their potential as early indicators of waterlogging stress. Our findings show that hyperspectral indices captured early physiological signals whereas RGB sensors only detected significant differences from Day 3 onwards, reflecting the lag between physiological stress and visible growth reduction. Our striking observation of such early significant changes in hyperspectral indices supports our AI results, where indices linked to pigment dynamics and canopy structure were identified as the top predictors of waterlogging stress. XAI highlighted three hyperspectral indices — MCARI1, SIPI and WATER1 — consistently across all algorithms as key indicators of waterlogging stress responses. Our results align with evidence showing that spectral indices reflecting pigment dynamics and water status are among the most sensitive early markers of abiotic stress. MCARI1 has been shown to respond strongly to chlorophyll-related changes associated with drought and water stress in Arabic coffee and maize, as demonstrated using multispectral and UAV-derived vegetation indices (Zhang *et al*., 2021; Da Silva *et al*., 2024), while SIPI captures very early shifts in leaf reflectance that reliably distinguish water-stressed from non-stressed plants in perennial grasses and olive plants, and more broadly reflects crop physiological status linked to pigment changes (Sun *et al*., 2014; Zhou *et al*., 2019; Katuwal *et al*., 2023). Likewise, WATER1 was highly informative, consistent with studies demonstrating that water-absorption-based indices closely track variation in leaf and canopy water content under water stress in crops such as grapevine, hibiscus, and spring wheat (Serrano *et al*., 2010; Shimada *et al*., 2012; Zununjan *et al*., 2024).

Overall, AI algorithms are well suited to capture the temporal complexity of waterlogging responses, particularly when supported by hyperspectral indices. The consistent importance of PSA, MCARI1, SIPI and WATER1 illustrates how XAI and its algorithms can pinpoint biologically meaningful features even in noisy phenomics’ datasets. Our insights highlight the value of XAI algorithms for interpreting dynamic and complex physiological responses occurring under abiotic stress.

### GWAS Interaction model reveals new associations between hyperspectral indices and stress tolerance

Our GWAS analysis detected 12 LD blocks as associated with traits contributing to waterlogging responses in three phases, highlighting two traits as particularly important. The first trait, WATER1, represents an early stress response with QTLs identified from days 2 to day 5. The second trait, PSRI, was found to be associated with the recovery phase. While both WATER1 and PSRI exhibited low heritability when calculated over the full 21-day experiment, this measure may not fully capture their genetic contributions due to their contrasting temporal dynamics. Our results corroborate previous studies that have demonstrated the effectiveness of these hyperspectral indices in detecting plant stress. For instance, (Behmann *et al*., 2014) reported that PSRI could detect drought stress in barley, while (Roy *et al*., 2023) showed the utility of leaf water vegetation indices in identifying susceptible genotypes in wheat under drought conditions. An absence of association was noted in the late stress stage (days 8-14). We hypothesise that this lack is due to substantial genomic activity occurring in the early stress phase as the plant begins to adapt, as well as heightened activity during recovery. By the late stress stage, the plant has predominantly returned to normal functioning.

In this work, we employed two distinct GWAS models: BLINK, which is particularly suitable for smaller datasets and has been successfully applied to this collection (Bernád *et al*., 2024), and an Interaction model introduced by Al-Tamimi *et al*. (2016). The Interaction model accounts for the interaction between treatment (control and waterlogged) and the genetic markers of interest (Al-Tamimi *et al*., 2016). The GWAS Interaction model has been shown to have a superior performance compared to a traditional MLM model (Al-Tamimi *et al*., 2016). As reported by Al-Tamimi *et al*. (2016), we found the same superior performance of the Interaction model compared to BLINK. Our results show that the Interaction model identified more QTLs and significant SNPs that remained consistent over several days, whereas BLINK was only capable of detecting significant association in one-off time points. Furthermore, we enhanced the accessibility of the Interaction model, which was previously implemented by Al-Tamimi *et al*. (2016) using ASReml, a proprietary software. Our implementation utilises base R, and the script has been made open access for the GWAS research community https://github.com/Walshj73/3D-QTLVis.

Reconciling research conducted in centimorgans (cM) with studies using base pairs (bp) presents a growing challenge. While the two metrics are comparable, precise conversion remains difficult due to their different foundations: cM measures recombination frequency during meiosis whereas bp provides a physical nucleotide count. These metrics serve distinct purposes: cM is used for mapping genetic linkage and linkage disequilibrium (LD), whereas bp offers a stable framework for gene annotation and cross-study integration. This shift from cM — still favoured by private-sector breeders — to bp reflects a broader move toward multi-omics integration. This challenging transition prompts a re-evaluation of how genomic data is utilised in modern breeding and genetic analysis. Hence, we found limited overlap of identified QTLs with the existing literature. The only point of congruence was identified with (Borrego-Benjumea *et al*., 2021), who reported a QTL associated with Spike, Kernel Weight and Grains per Plant linked to the candidate gene HORVU5Hr1G087730, which encodes the 13S globulin seed storage protein 2. This gene is located within the same LD block as our found H5-515. We hypothesise that the lack of shared QTL results can be attributed to earlier publications using cM distances, making meaningful comparisons to base pairs challenging. Furthermore, the limited overlap between our results and other GWAS studies can also be attributed to differing experimental conditions, selected growth stages, or phenotyping methods. For example, (Borrego-Benjumea *et al*., 2021) conducted their research under field conditions, with data collected at harvest, while (Manik *et al*., 2022) concentrated exclusively on root phenotyping and specifically adventitious root formation. Similarly, Luan *et al*., (2022) investigated waterlogging stress during germination.

Interestingly, our GWAS and XAI results pinpoint WATER1 as a major waterlogging stress predictor as well as significantly associated with nine LD blocks. Consistent with expectations, this hyperspectral index reflects variations in leaf and canopy water content, underscoring the key role of water dynamics in early waterlogging stress. However, GWAS and AI algorithms differed in highlighting key waterlogging traits. GWAS revealed genetic loci linked to WATER1 and PSRI, whereas AI algorithms highlighted PSA, SIPI, MCARI1, and WATER1 as top features. We may conclude that this divergence likely reflects the distinct objectives of the two approaches: AI seeks to identify the most robust phenotypic predictors of stress and recovery, whereas GWAS aims to uncover traits with a strong underlying genetic architecture. In other words, a trait’s utility as an early stress predictor does not necessarily imply a strong underlying genetic component. Furthermore, the GWAS models used (BLINK and Interaction) do not inherently account for the longitudinal nature of HTP data. To address this, we restricted our focus to QTLs that remained significant across multiple days, ensuring the robustness of the associations. Although our used GWAS models are not longitudinal, this multi-day filtering ensured that we effectively captured the temporal dynamics of waterlogging stress and adaptation.

### 3D-QTLVis highlights the dynamic genomic shifts through time

Visualising longitudinal GWAS results presents a significant challenge, yet it is a major factor for making sense of complex phenotyping data. Although numerous HTP tools exist for data generation, far fewer offer the advanced downstream capabilities required to analyse and visualise the high-dimensional datasets generated by HTP and GWAS. For instance, the IHUP platform integrates extraction and analysis but is largely tailored to Unmanned Aerial Vehicle (UAV) specific pipelines (B. Wang *et al*., 2024), HSI-PP focuses solely on hyperspectral data (ElManawy *et al*., 2022), and PlantCV, despite its flexibility, still requires substantial user expertise for advanced analytical workflows (Gehan *et al*., 2017; Sheng *et al*., 2023). Furthermore, conventional visualisation of GWAS results, such as Manhattan plots, are poorly suited to longitudinal datasets as they obscure dynamic temporal changes by presenting them as static snapshots. As genetic effects can appear, disappear or shift direction over time, recent studies have highlighted the need for visualisation frameworks that capture dynamic genetic architectures in time-resolved phenotyping data (Al-Tamimi *et al*., 2016; Agnew *et al*., 2024). To address this gap, we developed 3D-QTLVis, a simplified and accessible R tool for exploring longitudinal GWAS outputs in an interactive 3D space.

Our 3D-QTLVis tool incorporates several features to facilitate data exploration and enhance the understanding of complex longitudinal datasets. The 3D-QTLVis includes an interactive 3D Manhattan plot that allows users to intuitively navigate the data, zoom in on a specific SNP of interest, and examine its significance over time. It also includes a boxplot to visualise phenotypic variation and allelic differences under the tested conditions. To illustrate the utility of 3D-QTLVis, we used the LD block H5-515 as a representative example. The visualisation clearly captured the temporal evolution of this region, allowing us to identify early emerging peaks and track their distinct changes over time. From this analysis, two SNPs were particularly prominent, with significance values proximal to the threshold. Upon closer examination, and by clicking on the second window, the line graph revealed a low LOD score on day one, which sharply increased during days 2–5, followed by a gradual decline over the remainder of the experiment. The corresponding boxplot in window 3 showed clear allelic interactions with stress, providing valuable insights into the phenotypic variation driven by genetic differences. This integrated, interactive framework bridges genomic signals with time-series data, underscoring the value of 3D-QTLVis for understanding how genetic effects evolve throughout an experiment. Nevertheless, the performance of our tool was limited by the technical limitation of the R Shiny package to load the original HTP datasets and displaying the interactive 3D data. To enhance scalability, future iterations of the 3D-QTLVis should test alternative 3D libraries/packages in both R and Python. For example, PyVista, VisPy, or Mayavi may offer superior rendering capabilities compared to Matplotlib, enabling the interactive analysis of larger, more complex genomic datasets (Campagnola *et al*., 2025; Ramachandran and Varoquaux, 2011; Sullivan and Kaszynski, 2019).

Ultimately, 3D-QTLVis enabled us to pinpoint the dynamic genomic shifts in barley to waterlogging across the HTP experiment. The candidate genes identified here offer significant insights into the complex mechanisms underlying waterlogging tolerance in barley. The most significant candidate gene groups identified include those involved in oxidative stress, cell wall modification, and transcriptional regulation — pathways integral to the hypoxic response typical of waterlogging tolerance. We also highlight the potential of HTP to identify candidate genes that are linked to specific stress phases. For example, during the early stress response, candidate genes such as HORVU.MOREX.r3.5HG0428280 (Peroxidase), HORVU.MOREX.r3.5HG0429400, HORVU.MOREX.r3.5HG0429380 & HORVU.MOREX.r3.5HG0534650 (Glutaredoxin domain-containing protein), and HORVU.MOREX.r3.5HG0533860 (NAC domain-containing protein) play critical roles in managing oxidative stress and initiating stress signalling pathways, making them particularly relevant to an early stress phase (Mhamdi and Van Breusegem, 2018). Conversely, the recovery phase is characterised by genes such as HORVU.MOREX.r3.2HG 0202910 (Glutathione S-transferase), HORVU.MOREX.r3.2HG0203150 (Sucrose transporter), HORVU. MOREX.r3.2HG0203240 (WAT1-related protein), and HORVU.MOREX.r3.3HG0327 (Auxin efflux carrier component), which are involved in detoxification, energy distribution, and restoring physiological balance (Geldhof *et al*., 2022).

Notably, GWAS studies in barley under waterlogging stress have identified candidate genes belonging to the MYB transcription factor family (Luan *et al*., 2022; Manik *et al*., 2022; Borrego-Benjumea *et al*., 2021). In fact, MYB transcription factors have been extensively documented in barley responses to waterlogging. They are involved in aerenchyma formation (Sherwood et al., 2025), influence cell wall integrity and stress resilience (F. Wang *et al*., 2024), manage sugar metabolism (F. Wang *et al*., 2024), and promote survival mechanisms through anaerobic respiration and ROS management (Miricescu *et al*., 2023; F. Wang *et al*., 2024). The identification of MYB transcription factors across various stress phases suggests that they play a significant role in the waterlogging response, highlighting the need for further investigation into their functions and mechanisms.

In conclusion, our study combines HTP, explainable AI, and longitudinal GWAS, and provides a powerful framework for dissecting stress tolerance. Temporal imaging of barley under waterlogging used clustering analysis, revealing three distinct stress response phases, with PSA, WATER1, SIPI, and MCARI1 emerging as the most informative indicators of stress progression according to explainable AI. We found 236 SNPs across multiple LD blocks, exposing dynamic genetic architecture underpinning stress and recovery responses. We identified several candidate genes involved in oxidative stress defence, transcriptional regulation, auxin transport, and cell wall modification, with MYB transcription factors consistently implicated across phases — highlighting their central role in waterlogging adaptation. To support interpretation of longitudinal GWAS, we developed 3D-QTLVis, an interactive tool that visualises SNP–trait relationships across genomic and temporal dimensions. While future improvements in scalability are needed, this tool enables clearer identification of temporal QTL patterns. Collectively, our work provides valuable genetic and phenotypic resources for improving barley waterlogging tolerance and demonstrates the importance of integrating AI-driven phenotyping with temporal genomic analysis. This combined approach will be essential for the future development of robust predictive tools and accelerating breeding for resilience to abiotic stress.

## Supplementary data

Supplementary Fig. S1. Polynomial curves of the Relative Growth Rate (RGR) over 14 days of waterlogging followed by 7 days of recovery.

Supplementary Fig. S2. Linear correlation between the variables projected shoot area (px) and dry weights (g).

Supplementary Fig. S3. Polynomial curves of NDVI through 14 days of waterlogging followed by 7 days of recovery.

Supplementary Fig. S4. Polynomial curves of NDVI2 through 14 days of waterlogging followed by 7 days of recovery.

Supplementary Fig. S5. Polynomial curves of OSAVI through 14 days of waterlogging followed by 7 days of recovery.

Supplementary Fig. S6. Polynomial curves of PRI through 14 days of waterlogging followed by 7 days of recovery.

Supplementary Fig. S7. Polynomial curves of PSRI through 14 days of waterlogging followed by 7 days of recovery.

Supplementary Fig. S8. Correlation matrix between eleven imaging indices.

Supplementary Table S1. Date and number of accessions of the experimental runs carried out at the University of Picardie Jules Verne (UPJV).

Supplementary Table S2. Summary of the Hyperspectral Vegetation Indices, including abbreviation, full name, equation, and description of each index.

Supplementary Table S3. Heritability of eleven imaging phenotypic traits over 21 days under controlled and waterlogged conditions.

Supplementary Table S4. List of significant marker-trait associations detected by GWAS using BLINK and Interaction models under controlled conditions.

Supplementary Table S5. Summary of candidate genes containing significant markers associated with the traits evaluated under waterlogging treatment under controlled conditions, grouped according to functional annotation from GO terms.

## Acknowledgements

We thank Hervé Demailly, Gaëlle Mongelard, Stéphanie Guénin and Paul Ruel for technical assistance during the HTP experiments.

## Author contributions

SN, EM, and LG: conceptualization; EJ and VB: methodology; VB, JJW, EJ, and MK: formal analysis; VB, JJW, and EJ: investigation; NZ, KW, FF, LC, WBS, and JJW: software; VB and JJW: visualization; VB, JJW, and SN: writing — original draft; VB, JJW, SN, EM, EJ, LG, MK, and WBS: writing — review & editing; SN, EM, and LG: supervision. VB and JJW contributed equally to this work.

## Data availability

Genotypic data for the ExHIBiT core collection are available as described in Bernád *et al*. (2024). Summary statistics for all phenotypic data are provided in the supplementary data. Code used for running the GWAS interaction model in R and the 3D-QTLVis tool are available at https://github.com/Walshj73/3D-QTLVis.

