## Supplemental Figures for "Integrating longitudinal hyperspectral phenotyping with AI and GWAS to dissect barley waterlogging responses": Supp. Figures.docx

### *JXB* Supporting Information

The following Supporting Information is available for this article:

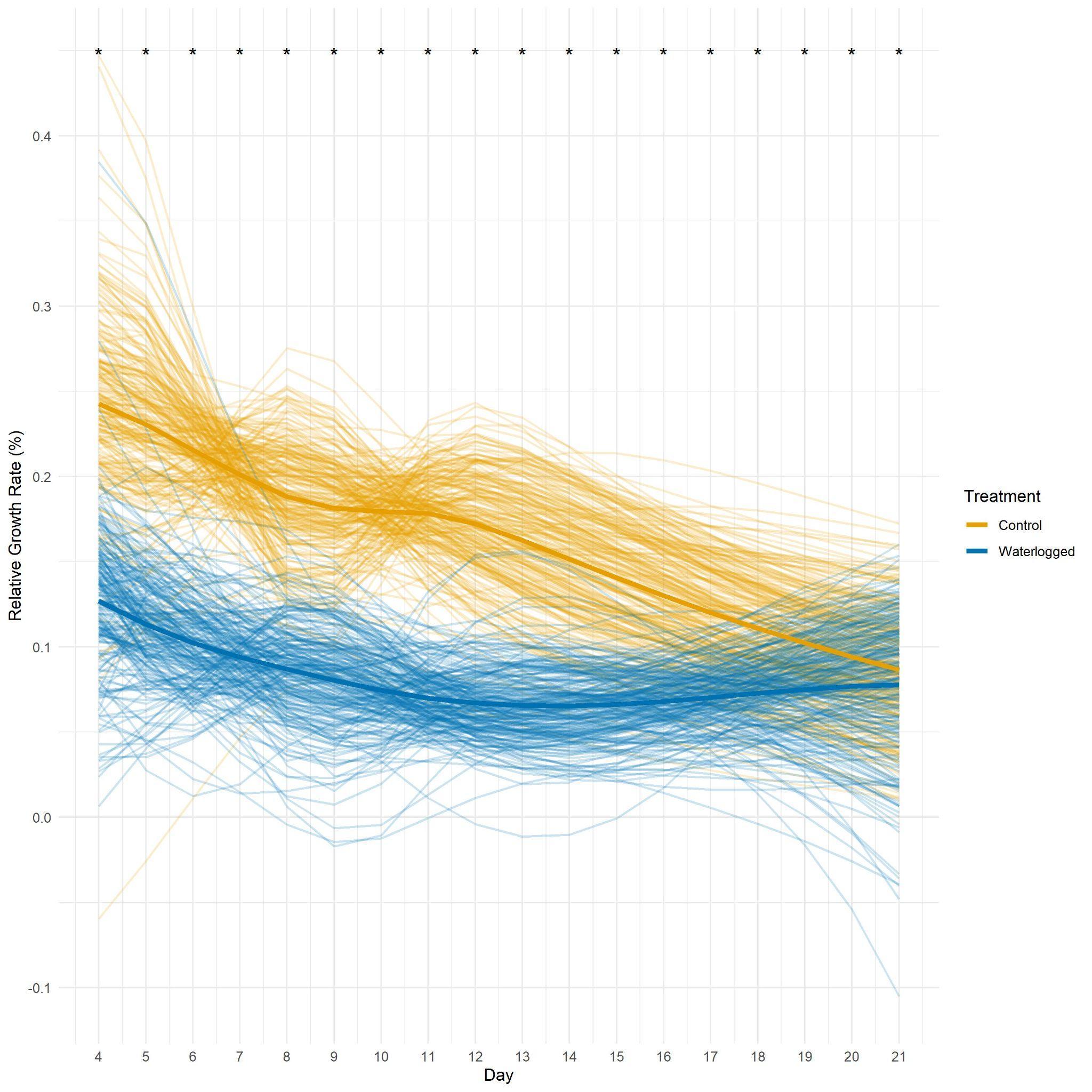

**Fig. S1** Fig S1. Polynomial curves of the Relative Growth Rate (RGR) over 14 days of waterlogging followed by 7 days of recovery. Yellow lines represent the control treatment, while blue lines represent the waterlogged treatment. Each thin line corresponds to an individual accession, and the bold lines depict the average values for each treatment. Black stars indicate statistically significant differences between control and waterlogged treatments on specific days, with significance determined at a p-value threshold of 0.05.

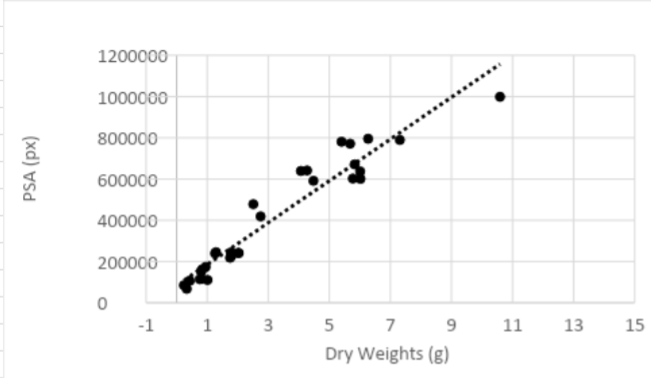

Fig S2. Linear correlation between the variables projected shoot area (px) and dry weights (g). Control and waterlogged plants of three different cultivars out of the core-collection were randomly chosen in every independent runs.

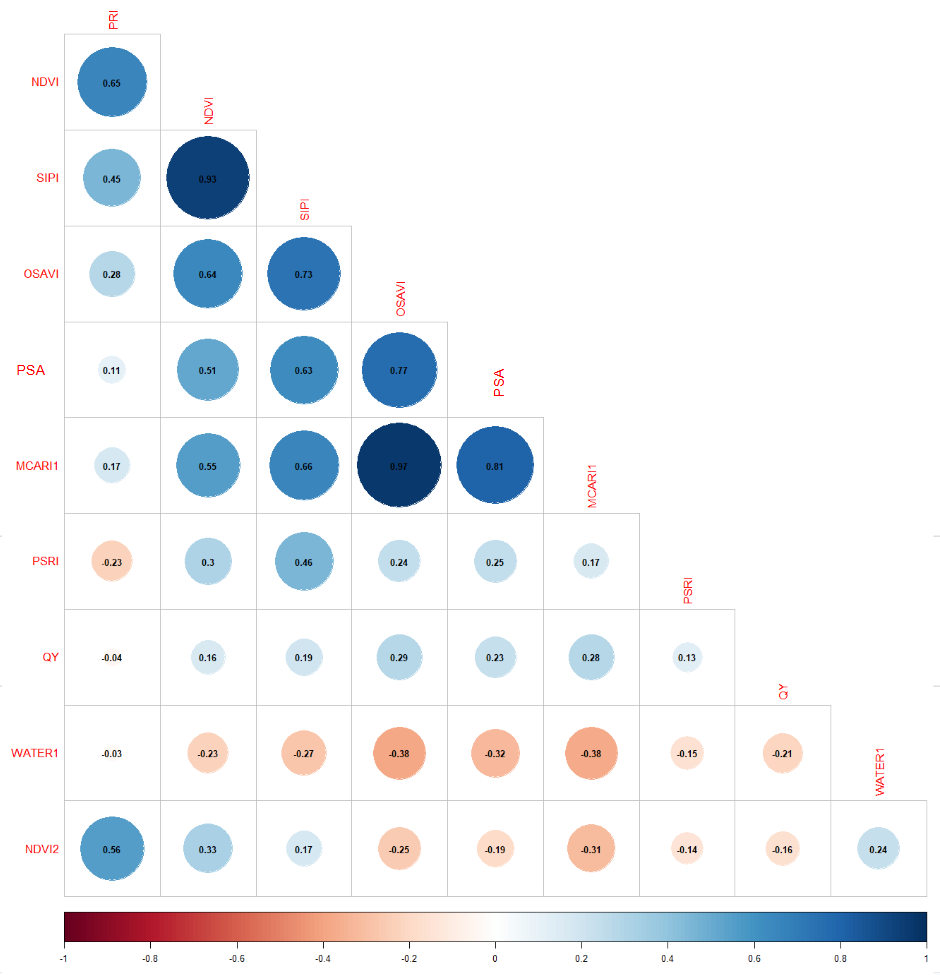
Fig S3. Correlation matrix between eleven imaging indices. Blue shows positive correlation and red negative correlation. Only significant plots are displayed.

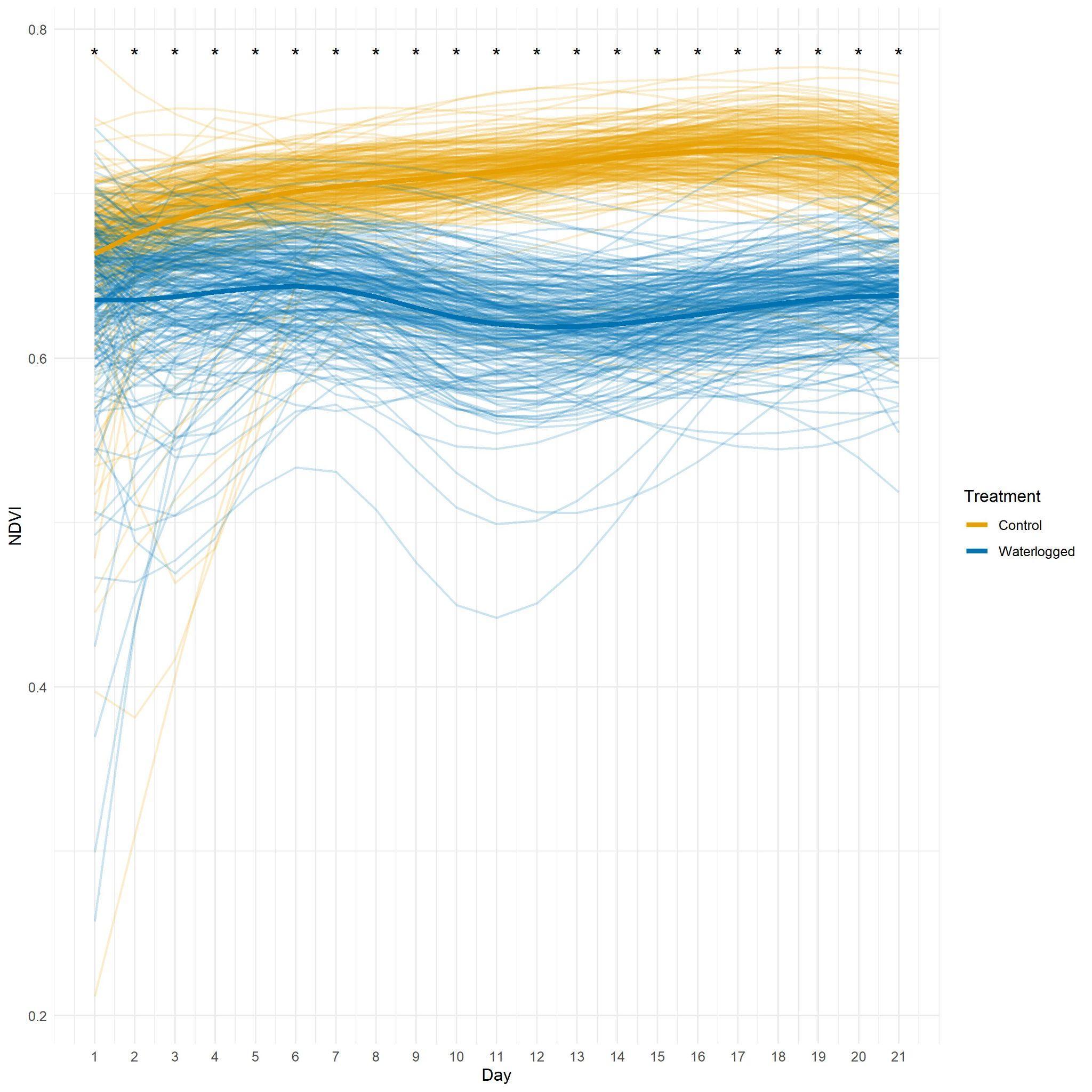

Fig S4. Polynomial curves of NDVI through 14 days of waterlogging followed by 7 days of recovery. Yellow lines represent the control treatment, while blue lines represent the waterlogged treatment. Each thin line corresponds to an individual accession, and the bold lines depict the average values for each treatment. Black stars indicate statistically significant differences between control and waterlogged treatments on specific days, with significance determined at a p-value threshold of 0.05

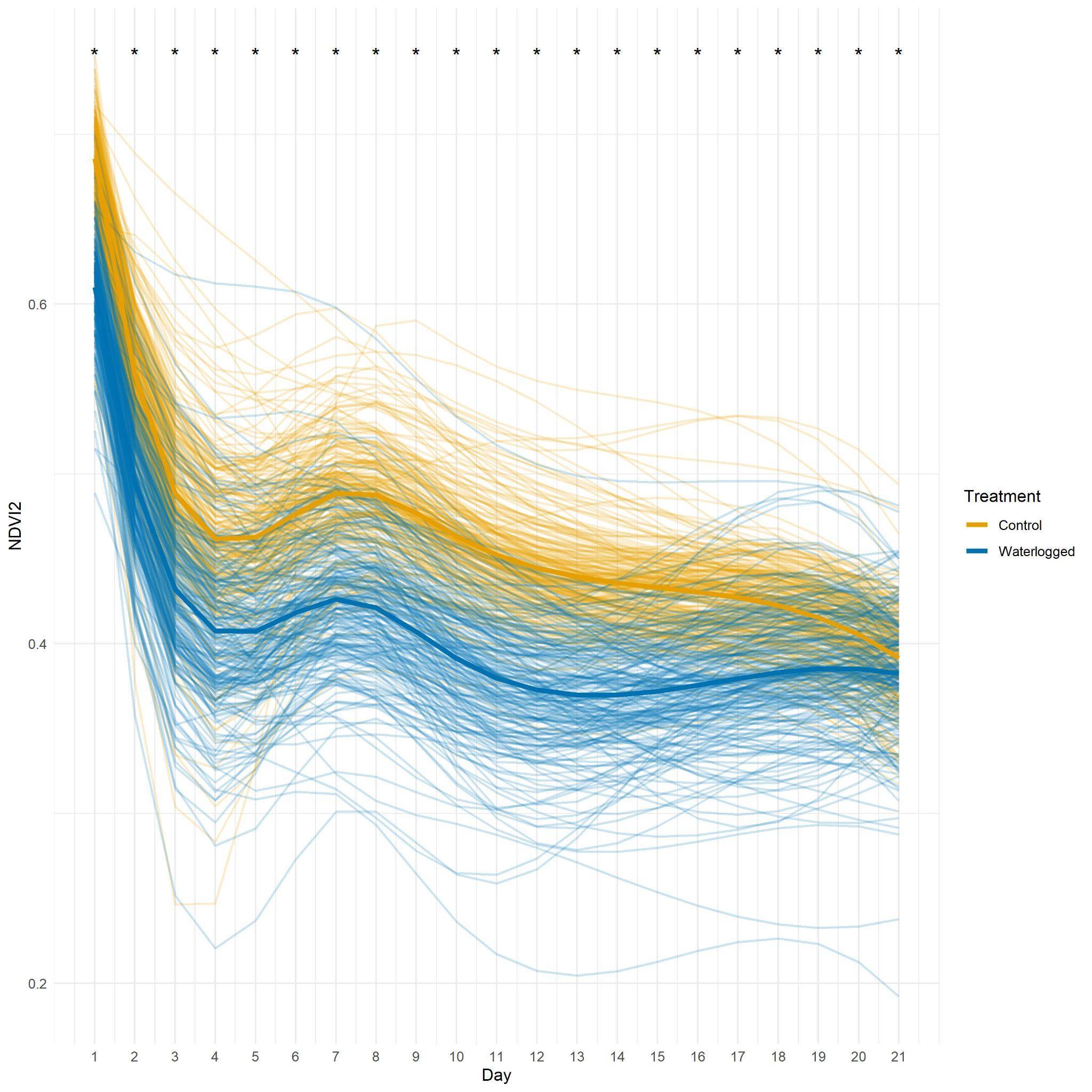

Fig S5. Polynomial curves of NDVI2 through 14 days of waterlogging followed by 7 days of recovery. Yellow lines represent the control treatment, while blue lines represent the waterlogged treatment. Each thin line corresponds to an individual accession, and the bold lines depict the average values for each treatment. Black stars indicate statistically significant differences between control and waterlogged treatments on specific days, with significance determined at a p-value threshold of 0.05.

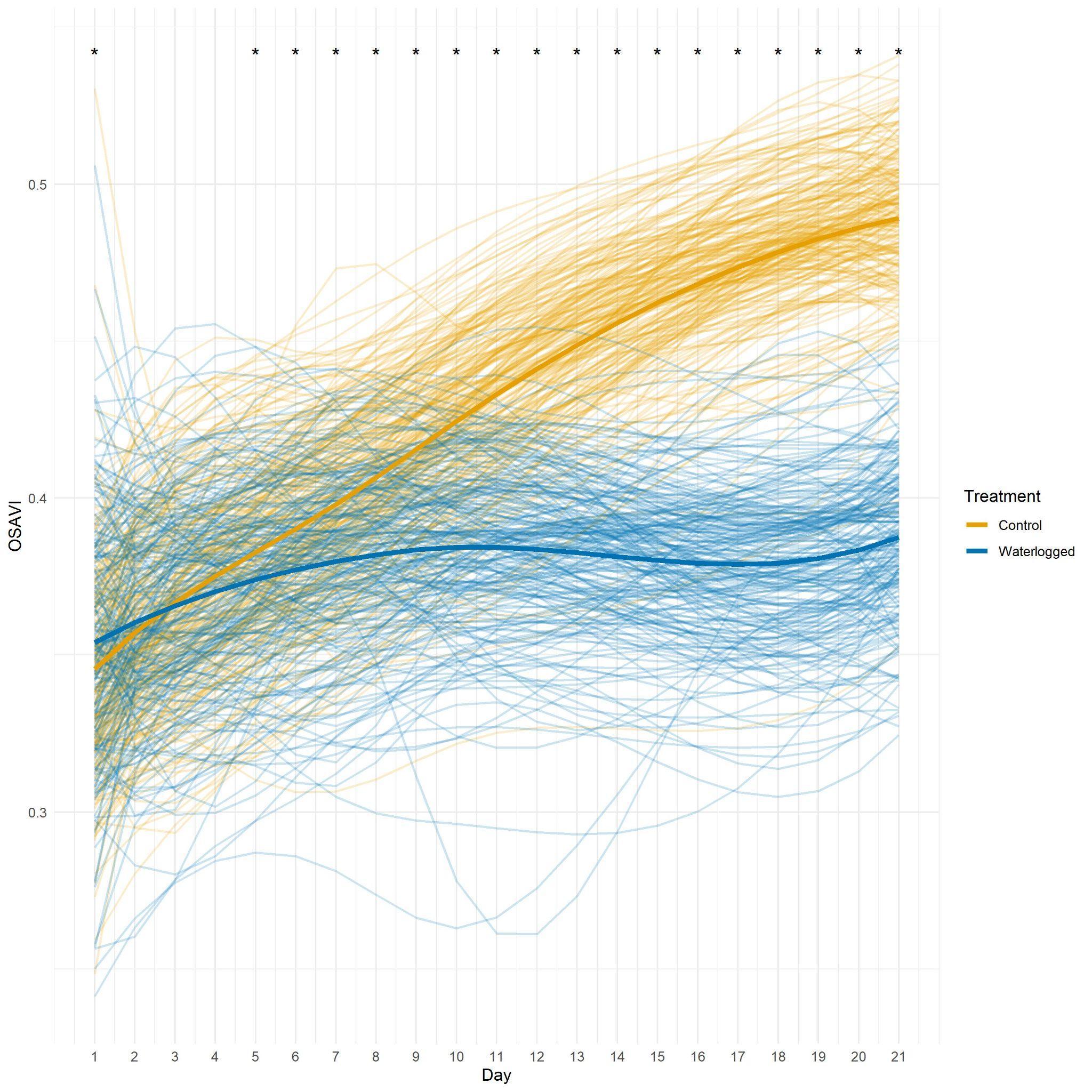

Fig S6. Polynomial curves of OSAVI through 14 days of waterlogging followed by 7 days of recovery. Yellow lines represent the control treatment, while blue lines represent the waterlogged treatment. Each thin line corresponds to an individual accession, and the bold lines depict the average values for each treatment. Black stars indicate statistically significant differences between control and waterlogged treatments on specific days, with significance determined at a p-value threshold of 0.05.

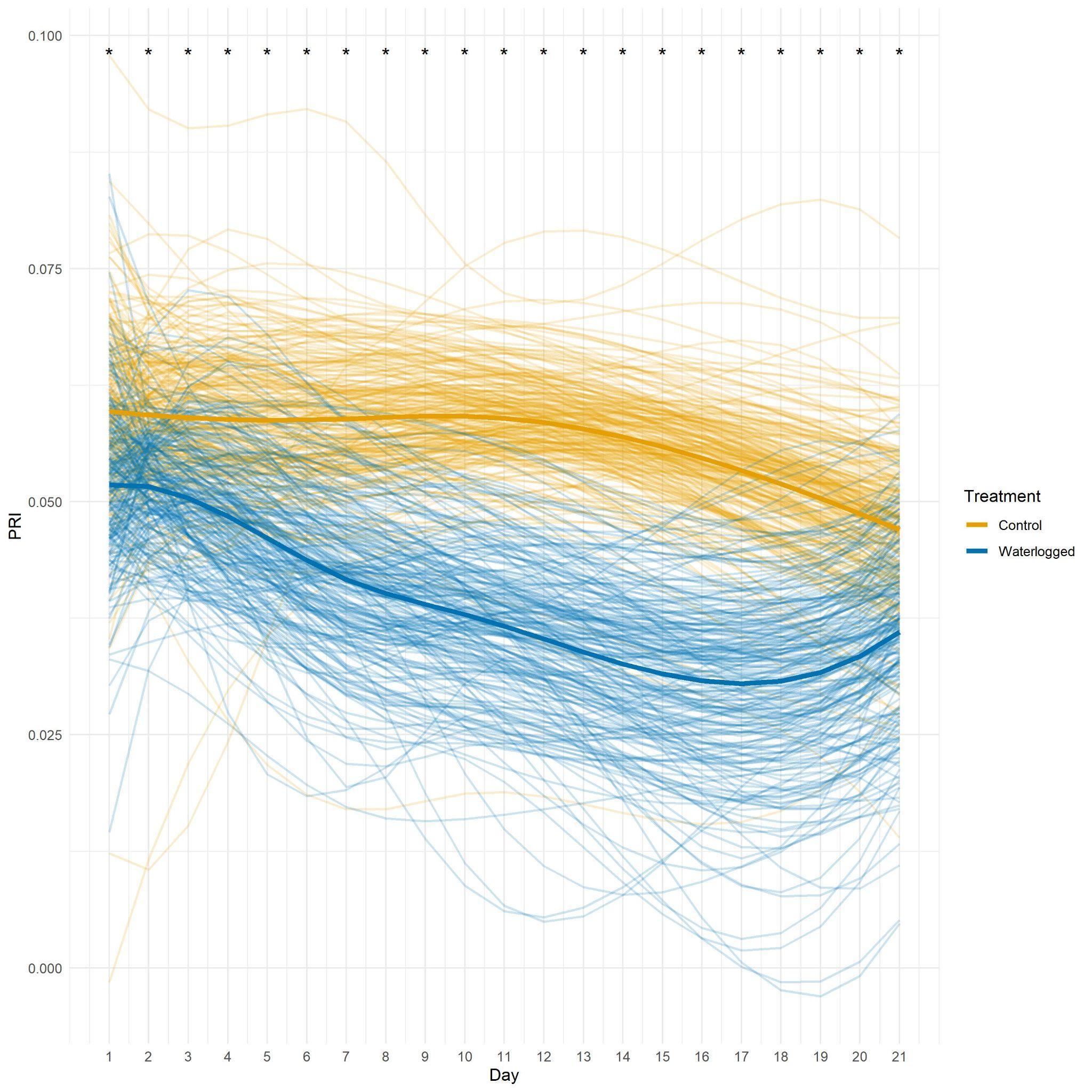

Fig S7. Polynomial curves of PRI through 14 days of waterlogging followed by 7 days of recovery. Yellow lines represent the control treatment, while blue lines represent the waterlogged treatment. Each thin line corresponds to an individual accession, and the bold lines depict the average values for each treatment. Black stars indicate statistically significant differences between control and waterlogged treatments on specific days, with significance determined at a p-value threshold of 0.05.

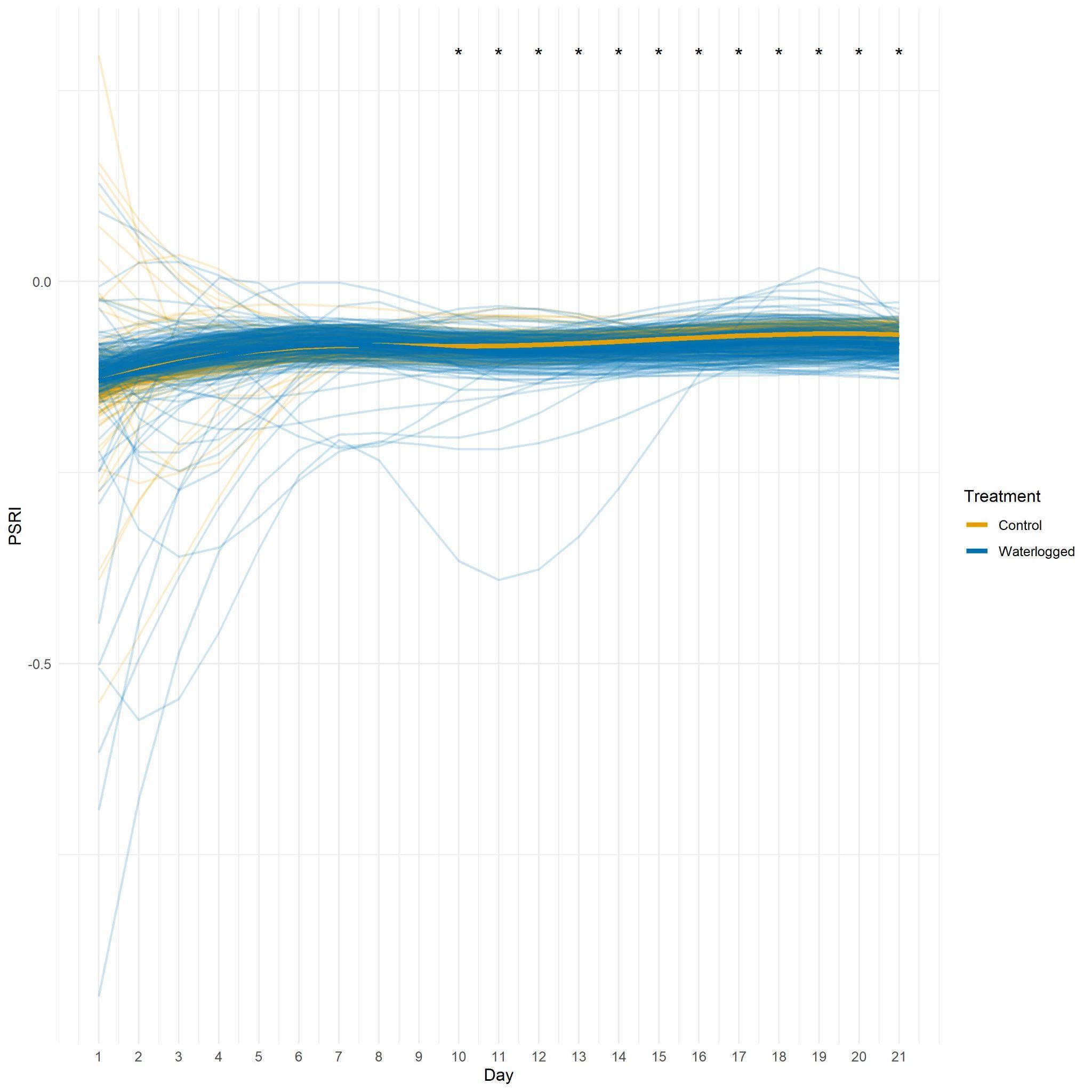

Fig S8. Polynomial curves of PSRI through 14 days of waterlogging followed by 7 days of recovery. Yellow lines represent the control treatment, while blue lines represent the waterlogged treatment. Each thin line corresponds to an individual accession, and the bold lines depict the average values for each treatment. Black stars indicate statistically significant differences between control and waterlogged treatments on specific days, with significance determined at a p-value threshold of 0.05.
